# Snails show their true colours: rate of evolutionary divergence in the shell colour polymorphism of *Cepaea nemoralis*

**DOI:** 10.64898/2026.09.14.749438

**Authors:** Govert Kwarten, Menno Schilthuizen

**Affiliations:** Institute of Biology Leiden, Leiden University, Leiden, Netherlands; Evolutionary Ecology Department, Naturalis Biodiversity Center, Leiden, Netherlands

## Abstract

The shell colour polymorphism in the land snail *Cepaea nemoralis* is maintained by environmentally variable light intensity and predation. Previous work showed that divergence in colour polymorphism between habitat pairs of open and closed areas increased with age of the habitat. However, that study was limited to a time span of 80 years (27 snail generations). Here, we investigate shell colour divergence over habitat pair ages of 95 to 325 years. We did find an increase in evolutionary divergence of colour polymorphism with increasing age, however equilibrium could not be determined. This is possibly due to the evolutionary effect of camouflage working in opposite direction to light intensity in certain locations, consequently clouding the results. Therefore, camouflage might also be an important factor to take into account in the search for the evolutionary equilibrium of colour polymorphism in *Cepaea nemoralis*.

## Introduction

Colour polymorphism is usually described as the presence of two or more clearly distinguishable, genetically determined colour morphs within a population, where the least common morph occurs too frequently to be explained by recurrent mutation (McKinnon & Pierotti, 2010). It is a widespread phenomenon that can be observed in many different species of organisms (Huxley, 1955). In evolutionary ecology, these polymorphisms make excellent study systems because many can be easily identified and scored in the field and are often affected by environmental selection, gene flow, and/or drift (McKinnon & Pierotti, 2010).

A well-known example of a species that exhibits colour polymorphism is the common European grove snail (*Cepaea nemoralis*). This snail species has three different shell ground colours: yellow, pink, and brown. Additionally, it exhibits different banding patterns and potential fusion of these bands (Schilthuizen, 2013). All of these traits are, almost entirely, genetically determined (Murray, 1975; Schilthuizen et al., 2020).

There are two main drivers for the natural selection of shell colour and banding pattern in *C. nemoralis*. The first of these is predation. This is habitat-dependent because if a morph blends in well with its surroundings, it is less likely to be preyed upon by visual diurnal predators. Predation, especially by the song thrush (*Turdus philomelos*), has been known to drive shell colouration towards habitat-specific camouflage (Cain & Sheppard, 1950; Lamotte, 1951). However, it also appears that the different *Cepaea nemoralis* morphs exhibit different behavioural strategies that possibly have evolved in response to predation. Unbanded morphs show a tendency to climb up the trees and avoid avian predators, while banded individuals tend to stay hidden on the ground (Rosin et al., 2018).

Secondly, temperature also seems to evolutionarily influence the frequencies of the different colour morphs in a population. This has to do with the thermoregulation of the snail; a lighter shell is better at reflecting sunlight than a darker shell. This is especially relevant because of ongoing climate change overall (Silvertown et al., 2011; Ożgo & Schilthuizen, 2012) and particularly in cities, where the temperature is often additionally elevated due to the urban heat island effect (Kerstes et al., 2019). Even without anthropogenic alteration of climate environments, heat as a selection still impacts morph frequencies in different environments, as in more open vegetation types the snails are exposed to much more direct sunlight than in closed vegetation types.

Besides the presence of its shell colour polymorphism, there are other reasons why *C. nemoralis* is a very suitable species for evolutionary ecology research. Firstly, as its common name suggests, this snail is quite abundant in large parts of Europe (including the Netherlands), and even in North America, where it has been introduced (Reed, 1964; Richards & Murray, 1975) which should make it easy to find numerous individuals in the field. Secondly, the shells are large, and the colour morphs are easy to recognise and score in the field, even in dead in individuals, juveniles, and shell fragments. Furthermore, the selection pressures acting upon this phenomenon are quite well known, which allows for targeted experimental and field studies.

In the present study, we aim to investigate the rate of evolutionary divergence of colour polymorphism across different habitats of different ages. Earlier research, making use of known ages of reclaimed land in The Netherlands, showed that at the oldest investigated age, approximately 80 years, which corresponds to 27 *Cepaea* generations, divergence between mean shell darkness for open and closed habitats was greatest (Schilthuizen, 2013). According to a model by Cook (1998), this divergence is expected to reach an equilibrium after approximately 300 years, or 100 snail generations. Here, we extend the temporal range to more than 300 years, to test the predictions from Cook’s model.

## Materials and methods

Shell colour polymorphism of *C. nemoralis* was scored using the same method as used in Schilthuizen (2013). The ground colour was marked as yellow (Y), brown (B) or pink (P), combined with a 5-digit code to indicate the banding pattern. Parentheses were used to indicate band fusions. Although in theory 3 × 5! = 360 colour morphs are possible, four banding morphs were most prevalent in the field: unbanded (00000), mid-banded (00300), triple-banded (00345), and five-banded (12345). Therefore, with three different ground colours, in practice, only 12 morphs were the most prevalent components of the shell colour polymorphisms. However, in rare cases other morphs can be found as well.

### Selection of research locations

We selected sample locations across the Netherlands on a scale ranging from the oldest around 300 years, to the youngest around 80 years. To connect our results to the earlier results by Schilthuizen (2013), we would have preferred choosing areas of reclaimed land of the appropriate ages. However, the use of reclaimed land for the purpose of this research does encounter problems, namely that older areas of reclaimed land, before high-quality dykes were developed, have been flooded frequently, potentially wiping out snail populations one or more times during their existence. Also, afforestation in these areas is probably from much more recent dates than the ages of the reclaimed land. Therefore, instead, we chose estates of known dates of establishment.

We selected sample locations from 3 different age categories: c. 1930, c. 1850, and c. 1700. To align with the previous research by Schilthuizen (2013), we also resampled the oldest of his areas of reclaimed land, the Wieringermeer, drained in 1930.

To select suitable research locations, we used a combination of satellite images to find potentially suitable habitats and the citizen science platform waarneming.nl to confirm the presence of *C. nemoralis*.

### Sampling method

For each research location, 1 habitat pair per research location was realized. Based on the known dispersal rate of *Cepaea* ranging from 5 to 10 m annually (Cameron, 2001), we attempted to preserve a minimum inter-habitat distance of 200 m. For each sample location, the aim was to collect at least 150 live, adult individuals. The sample locations were evaluated for the five most abundant tree and herb species and the type of ground coverage to increase insight into the overall characteristics of a specific habitat and allow for simple comparisons between habitats. All locations, their age categories, coordinates, and inter-habitat distances can be found in Table 1.

**Table 1:**
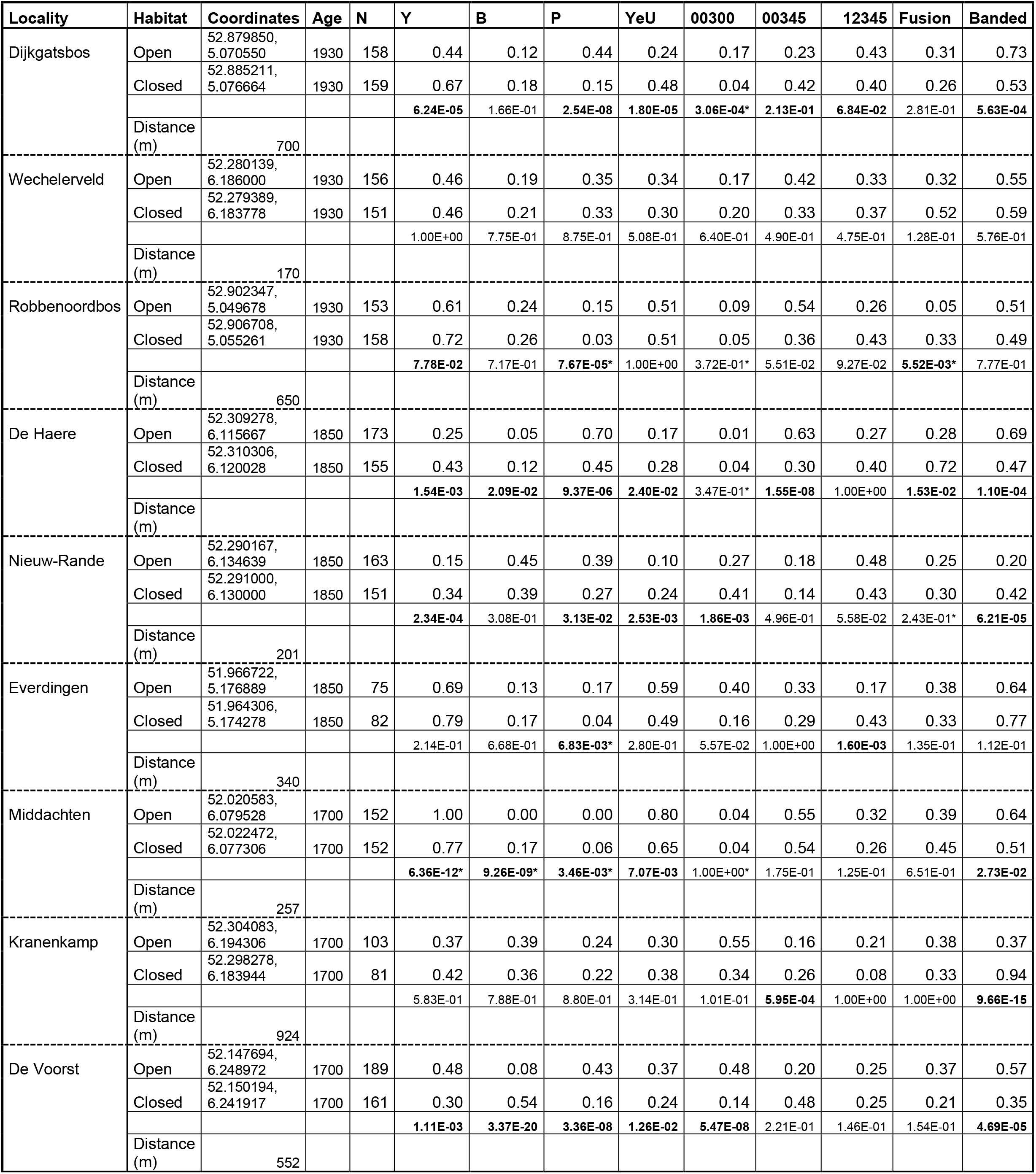
Showcases all 9 locations with coordinates and distance between sample locations. For the morph proportions, bold signals statistical significance (p<0.05). For marked p-values (*) a Fisher test was used instead.

| Locality | Habitat | Coordinates | Age | N | Y | B | P | YeU | 00300 | 00345 | 12345 | Fusion | Banded |
| --- | --- | --- | --- | --- | --- | --- | --- | --- | --- | --- | --- | --- | --- |
| Dijkgatsbos | Open | 52.879850,<br>5.070550 | 1930 | 158 | 0.44 | 0.12 | 0.44 | 0.24 | 0.17 | 0.23 | 0.43 | 0.31 | 0.73 |
|  | Closed | 52.885211,<br>5.076664 | 1930 | 159 | 0.67 | 0.18 | 0.15 | 0.48 | 0.04 | 0.42 | 0.40 | 0.26 | 0.53 |
|  |  |  |  |  | **6.24E-05** | 1.66E-01 | **2.54E-08** | **1.80E-05** | **3.06E-04\*** | **2.13E-01** | **6.84E-02** | 2.81E-01 | **5.63E-04** |
|  | Distance (m) | 700 |  |  |  |  |  |  |  |  |  |  |  |
| Wechelerveld | Open | 52.280139,<br>6.186000 | 1930 | 156 | 0.46 | 0.19 | 0.35 | 0.34 | 0.17 | 0.42 | 0.33 | 0.32 | 0.55 |
|  | Closed | 52.279389,<br>6.183778 | 1930 | 151 | 0.46 | 0.21 | 0.33 | 0.30 | 0.20 | 0.33 | 0.37 | 0.52 | 0.59 |
|  |  |  |  |  | 1.00E+00 | 7.75E-01 | 8.75E-01 | 5.08E-01 | 6.40E-01 | 4.90E-01 | 4.75E-01 | 1.28E-01 | 5.76E-01 |
|  | Distance (m) | 170 |  |  |  |  |  |  |  |  |  |  |  |
| Robbenoordbos | Open | 52.902347,<br>5.049678 | 1930 | 153 | 0.61 | 0.24 | 0.15 | 0.51 | 0.09 | 0.54 | 0.26 | 0.05 | 0.51 |
|  | Closed | 52.906708,<br>5.055261 | 1930 | 158 | 0.72 | 0.26 | 0.03 | 0.51 | 0.05 | 0.36 | 0.43 | 0.33 | 0.49 |
|  |  |  |  |  | **7.78E-02** | 7.17E-01 | **7.67E-05\*** | 1.00E+00 | 3.72E-01* | 5.51E-02 | 9.27E-02 | **5.52E-03\*** | 7.77E-01 |
|  | Distance (m) | 650 |  |  |  |  |  |  |  |  |  |  |  |
| De Haere | Open | 52.309278,<br>6.115667 | 1850 | 173 | 0.25 | 0.05 | 0.70 | 0.17 | 0.01 | 0.63 | 0.27 | 0.28 | 0.69 |
|  | Closed | 52.310306,<br>6.120028 | 1850 | 155 | 0.43 | 0.12 | 0.45 | 0.28 | 0.04 | 0.30 | 0.40 | 0.72 | 0.47 |
|  |  |  |  |  | **1.54E-03** | **2.09E-02** | **9.37E-06** | **2.40E-02** | 3.47E-01* | **1.55E-08** | 1.00E+00 | **1.53E-02** | **1.10E-04** |
|  | Distance (m) |  |  |  |  |  |  |  |  |  |  |  |  |
| Nieuw-Rande | Open | 52.290167,<br>6.134639 | 1850 | 163 | 0.15 | 0.45 | 0.39 | 0.10 | 0.27 | 0.18 | 0.48 | 0.25 | 0.20 |
|  | Closed | 52.291000,<br>6.130000 | 1850 | 151 | 0.34 | 0.39 | 0.27 | 0.24 | 0.41 | 0.14 | 0.43 | 0.30 | 0.42 |
|  |  |  |  |  | **2.34E-04** | 3.08E-01 | **3.13E-02** | **2.53E-03** | **1.86E-03** | 4.96E-01 | 5.58E-02 | 2.43E-01* | **6.21E-05** |
|  | Distance (m) | 201 |  |  |  |  |  |  |  |  |  |  |  |
| Everdingen | Open | 51.966722,<br>5.176889 | 1850 | 75 | 0.69 | 0.13 | 0.17 | 0.59 | 0.40 | 0.33 | 0.17 | 0.38 | 0.64 |
|  | Closed | 51.964306,<br>5.174278 | 1850 | 82 | 0.79 | 0.17 | 0.04 | 0.49 | 0.16 | 0.29 | 0.43 | 0.33 | 0.77 |
|  |  |  |  |  | 2.14E-01 | 6.68E-01 | **6.83E-03\*** | 2.80E-01 | 5.57E-02 | 1.00E+00 | **1.60E-03** | 1.35E-01 | 1.12E-01 |
|  | Distance (m) | 340 |  |  |  |  |  |  |  |  |  |  |  |
| Middachten | Open | 52.020583,<br>6.079528 | 1700 | 152 | 1.00 | 0.00 | 0.00 | 0.80 | 0.04 | 0.55 | 0.32 | 0.39 | 0.64 |
|  | Closed | 52.022472,<br>6.077306 | 1700 | 152 | 0.77 | 0.17 | 0.06 | 0.65 | 0.04 | 0.54 | 0.26 | 0.45 | 0.51 |
|  |  |  |  |  | **6.36E-12\*** | **9.26E-09\*** | **3.46E-03\*** | **7.07E-03** | 1.00E+00* | 1.75E-01 | 1.25E-01 | 6.51E-01 | **2.73E-02** |
|  | Distance (m) | 257 |  |  |  |  |  |  |  |  |  |  |  |
| Kranenkamp | Open | 52.304083,<br>6.194306 | 1700 | 103 | 0.37 | 0.39 | 0.24 | 0.30 | 0.55 | 0.16 | 0.21 | 0.38 | 0.37 |
|  | Closed | 52.298278,<br>6.183944 | 1700 | 81 | 0.42 | 0.36 | 0.22 | 0.38 | 0.34 | 0.26 | 0.08 | 0.33 | 0.94 |
|  |  |  |  |  | 5.83E-01 | 7.88E-01 | 8.80E-01 | 3.14E-01 | 1.01E-01 | **5.95E-04** | 1.00E+00 | 1.00E+00 | **9.66E-15** |
|  | Distance (m) | 924 |  |  |  |  |  |  |  |  |  |  |  |
| De Voorst | Open | 52.147694,<br>6.248972 | 1700 | 189 | 0.48 | 0.08 | 0.43 | 0.37 | 0.48 | 0.20 | 0.25 | 0.37 | 0.57 |
|  | Closed | 52.150194,<br>6.241917 | 1700 | 161 | 0.30 | 0.54 | 0.16 | 0.24 | 0.14 | 0.48 | 0.25 | 0.21 | 0.35 |
|  |  |  |  |  | **1.11E-03** | **3.37E-20** | **3.36E-08** | **1.26E-02** | **5.47E-08** | 2.21E-01 | 1.46E-01 | 1.54E-01 | **4.69E-05** |
|  | Distance (m) | 552 |  |  |  |  |  |  |  |  |  |  |  |

### Analyses

#### Scoring system for *Cepaea nemoralis*

For each population, we calculated the proportions of yellow (Y), brown (B), and pink (P); YeU (yellow effectively unbanded, i.e., any Y00XXX morph, where ‘X’ stands for either absence or presence of a band); mid-banded (X00300), triple-banded (X00345), and five-banded (X12345). Finally, the number of five-banded with at least one fusion and the total number of morphs with at least one fusion were determined. Any potentially significant differences between these proportions were assessed with chi-square tests. For proportions with counts <5, a Fisher test was used instead. Investigation of a potential increase of the inter-habitat differences with increasing age per morph category was performed using a Pearson correlation test, while an overall effect of age was estimated using a linear mixed model (R package lme4). Subsequent p-values were generated using the R package lmerTest.

For all snails, a darkness score was calculated, following the method of Schilthuizen (2013). To obtain this score, the data from Heath (1975) were used, where each colour, band, and band fusion will affect the albedo of the shell, and the darkness score represents the number of degrees (°C) added to the internal body temperature of the snail inside. Yellow was used as a baseline with a value of 0. The other 2 colour morphs, pink and brown, obtain +0.3 and +0.6 to this base value, respectively. For each band that a shell has, we added +0.07, and for every band fusion +0.03. This combined value comprises the darkness score. To assess significance between two sites, a Kruskal-Wallis Rank Sum test was used. This test was used because after testing the data using a Shapiro test, the assumption of the normality of the data appeared to be violated. A Levene test revealed the variances to be unequal.

Two locations in the Wieringermeer (Robbenoordbos and Dijkgatsbos) sampled by Schilthuizen (2013) in 2011, were resampled during this research and used to assess population differences over this 14-year interval.

Finally, to test for the potential influence of camouflage, a multivariate test was performed. Using a redundancy analysis (RDA) from the R package Vegan the 3 most abundant plant species were correlated to the morph proportions found in the different sample locations.

## Results

In most research locations, we found >150 snails. Only in Fort Everdingen (75 individuals in the open habitat and 82 in closed) and Kranenkamp (with 103 and 81 snails in open and closed, respectively), we did not achieve this lower limit. A total of 2572 *C. nemoralis* were sampled. In 7 locations, there was a significant difference between the proportion of pink morphs found between the habitat pairs. In 6 of these habitat pairs, pink morphs were more prevalent in the open areas. The difference in proportions of yellow morphs between habitats were significant in 6 locations, where in 4 locations yellow morphs were more prevalent in the open habitats. Contrary, brown morph proportions only differed in 3 locations where higher prevalence was always found in the closed areas. Table 1 clearly shows no increase in the total number of statistical differences between morph proportions per location with increasing age.

When all banded morphs are compared directly in unison, this morph frequency does show a surprising difference in the estate Kranenkamp, where 94% of all found snails were banded. Also, apart from Wechelerveld, Robbenoordbos, and Everdingen, there is a significant difference in the proportions of banded versus unbanded individuals between different habitats. However, a clear direction for an increase in bandedness in either shaded or open habitats cannot be observed as 4 locations have more banded morphs in the open half of the habitat pair versus 2 where this is the case in the closed area. The same lack of a distinct pattern can also be seen in the remaining proportions.

Furthermore, to look whether any differences in morph proportions increased with habitat age, all morph categories were graphed against time, figure 1. However, after using Pearson correlation tests to run a line through all the points per morph category, all 8 correlations came back not significant. To inspect a potential overall effect of age a linear mixed model corrected for repeated observations within locations was used, which estimated a not significant p-value of 0.5.

**Figure 1:**
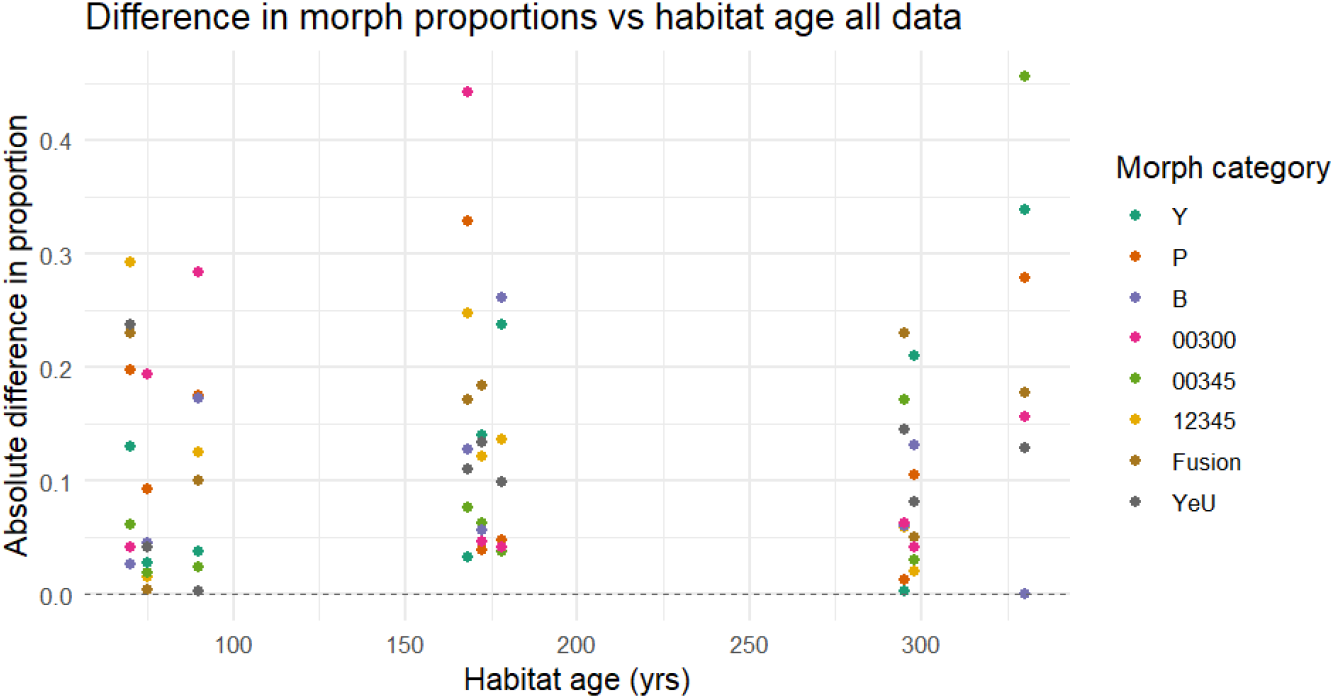
graph showing the absolute differences between morph frequencies per habitat pair plotted against time

The second step were the darkness scores. Out of the 9 habitat pairs, only 3 were not significantly different: Robbenoordbos, Wechelerveld and Kranenkamp. Out of the 6 statistically different locations, in 3 the mean darkness was greater in the open habitat, which was unexpected. See figure 2.

**Figure 2:**
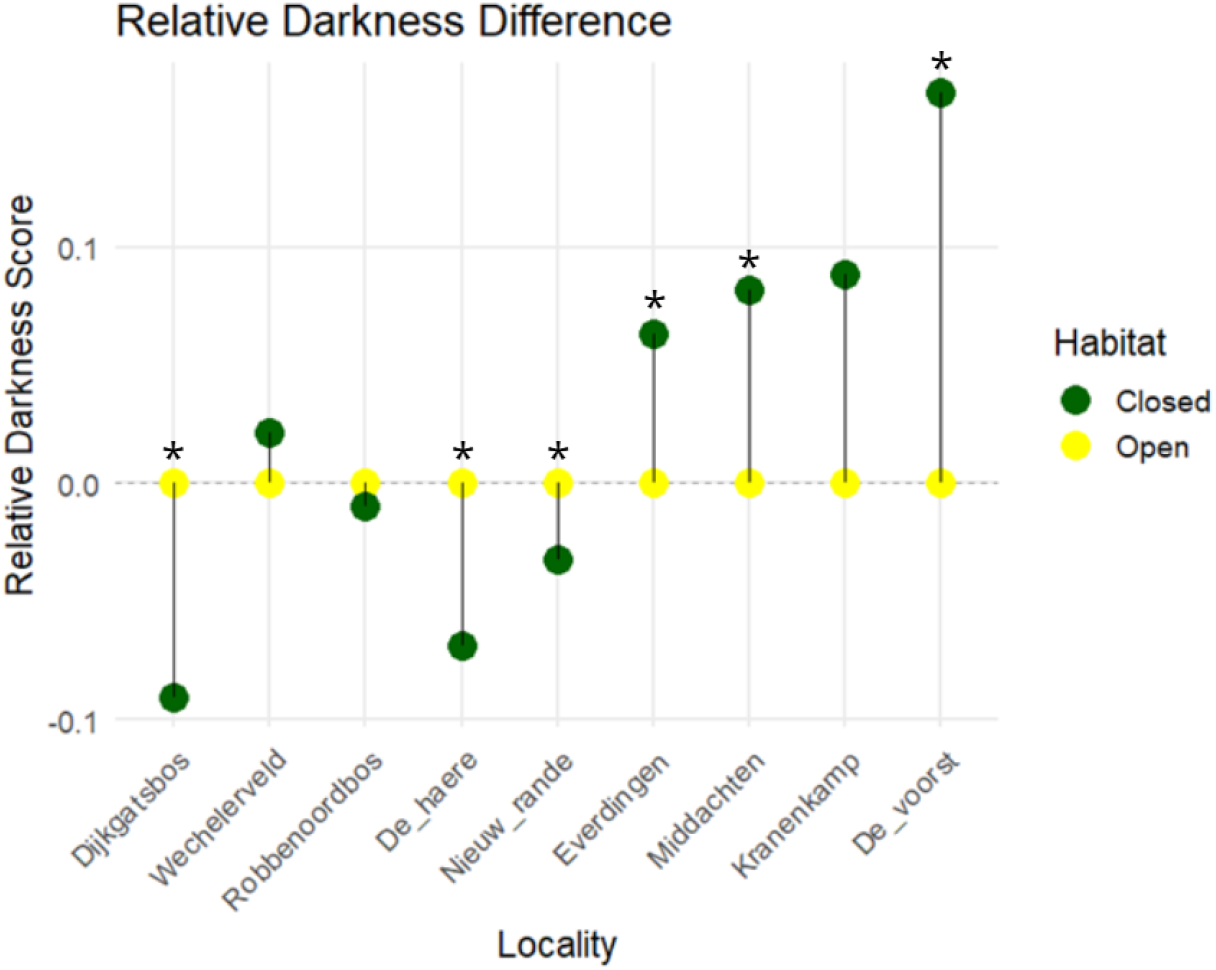
Graph showing the relative darkness score (closed-open) per habitat pair. * signals statistical significance (p<0.05).

Also, when plotting the absolute differences of these darkness scores per habitat pair against age and fitting a linear model through these values, there was a significant increase in darkness scores over time (p=0.03) (figure 3).

**Figure 3:**
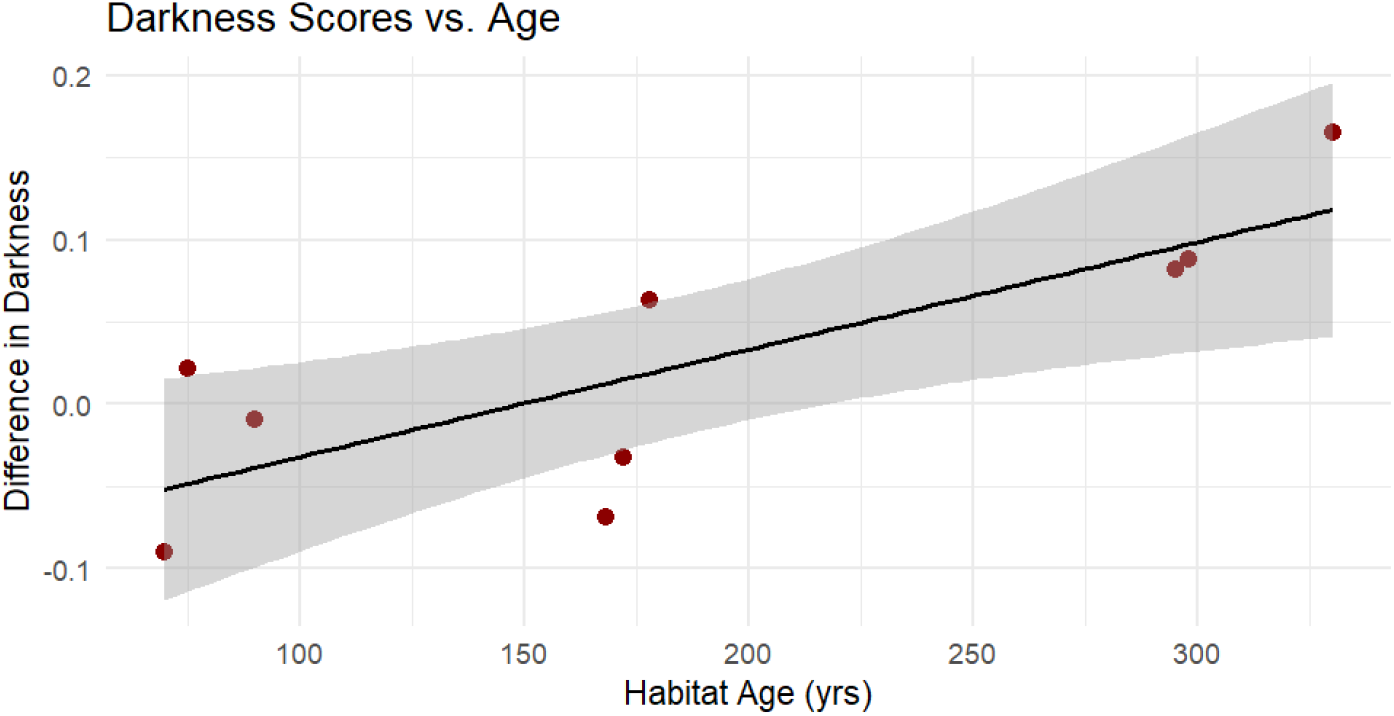
Differences in darkness scores against age.

Additionally, as we resampled two locations in the Wieringermeer (Robbenoordbos and Dijkgatsbos) of Schilthuizen (2013), we wished to check whether the results were similar to 14 years ago. Out of the 32 morph frequency comparisons between the new and old data, six comparisons were significantly different. For Dijkgatsbos (open) the 00345 and fusion categories were different. The other differences were in Robbenoordbos (closed) (Y, B, 00345, and YeU). The data from the other locations were not significantly different from the previous data, see table 2.

**Table 2:**
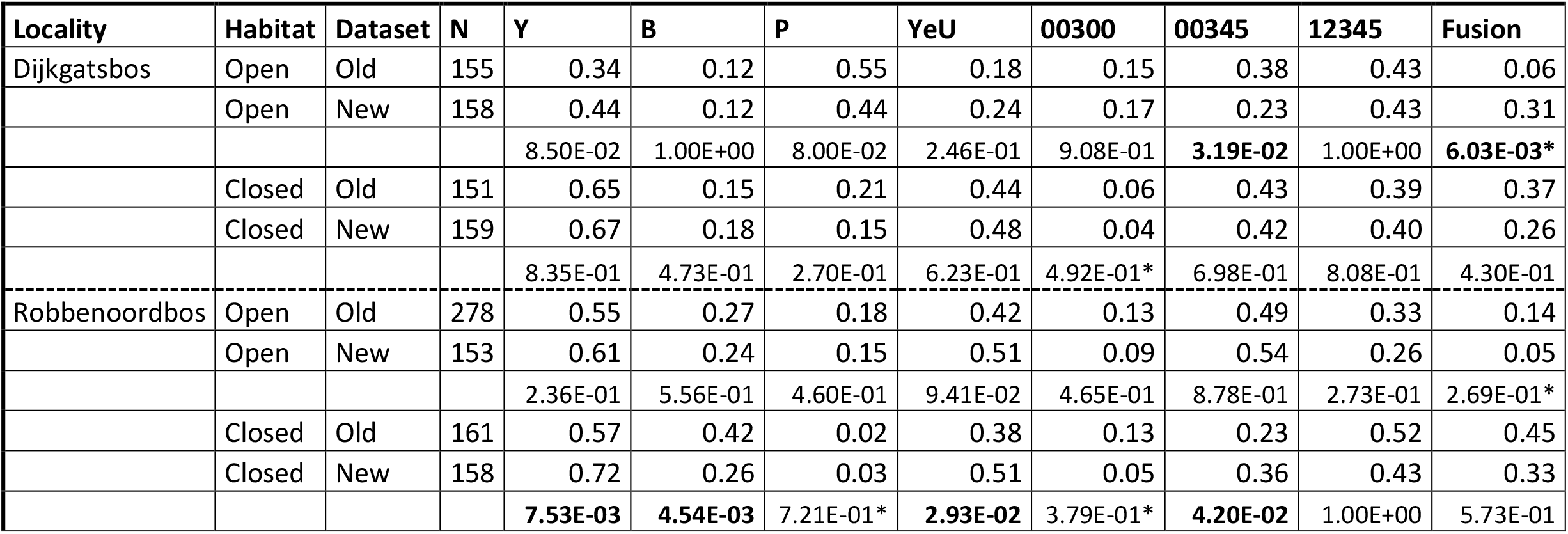
Shows morph proportions of the 2 resampled locations, accompanied with p-values. Bold signals statistical significance (p<0.05). For marked p-values (*) a Fisher test was used instead.

| Locality | Habitat | Dataset | N | Y | B | P | YeU | 00300 | 00345 | 12345 | Fusion |
| --- | --- | --- | --- | --- | --- | --- | --- | --- | --- | --- | --- |
| Dijkgatsbos | Open | Old | 155 | 0.34 | 0.12 | 0.55 | 0.18 | 0.15 | 0.38 | 0.43 | 0.06 |
|  | Open | New | 158 | 0.44 | 0.12 | 0.44 | 0.24 | 0.17 | 0.23 | 0.43 | 0.31 |
|  |  |  |  | 8.50E-02 | 1.00E+00 | 8.00E-02 | 2.46E-01 | 9.08E-01 | **3.19E-02** | 1.00E+00 | **6.03E-03\*** |
|  | Closed | Old | 151 | 0.65 | 0.15 | 0.21 | 0.44 | 0.06 | 0.43 | 0.39 | 0.37 |
|  | Closed | New | 159 | 0.67 | 0.18 | 0.15 | 0.48 | 0.04 | 0.42 | 0.40 | 0.26 |
|  |  |  |  | 8.35E-01 | 4.73E-01 | 2.70E-01 | 6.23E-01 | 4.92E-01* | 6.98E-01 | 8.08E-01 | 4.30E-01 |
| Robbenoordbos | Open | Old | 278 | 0.55 | 0.27 | 0.18 | 0.42 | 0.13 | 0.49 | 0.33 | 0.14 |
|  | Open | New | 153 | 0.61 | 0.24 | 0.15 | 0.51 | 0.09 | 0.54 | 0.26 | 0.05 |
|  |  |  |  | 2.36E-01 | 5.56E-01 | 4.60E-01 | 9.41E-02 | 4.65E-01 | 8.78E-01 | 2.73E-01 | 2.69E-01* |
|  | Closed | Old | 161 | 0.57 | 0.42 | 0.02 | 0.38 | 0.13 | 0.23 | 0.52 | 0.45 |
|  | Closed | New | 158 | 0.72 | 0.26 | 0.03 | 0.51 | 0.05 | 0.36 | 0.43 | 0.33 |
|  |  |  |  | **7.53E-03** | **4.54E-03** | 7.21E-01* | **2.93E-02** | 3.79E-01* | **4.20E-02** | 1.00E+00 | 5.73E-01 |

This pattern was sustained upon investigation of the darkness scores using a Kruskal-Wallis test, as the only significant difference was found between the shaded Robbenoordbos habitats (p=0.0053). The populations in the other locations remained unchanged in the past 14 years.

Finally, a multivariate analysis was carried out to gain insight into the effects of vegetation within habitats. The limited number of research locations and habitat pairs ensured the need for simplifying the model to only include three plant species: reeds (*Phragmites*), blackberries (*Rubus*), and nettle (*Urtica*). Figure 4 shows that all three plant species affect the data in different ways, with reeds correlating with yellow morphs while blackberries and nettle correlated with darker shell morphs. Because the habitat pairs are not independent, this analysis cannot be statistically verified. Still, this multivariate analysis does give an indication of the effect of certain plant species on morph frequency configurations.

**Figure 4:**
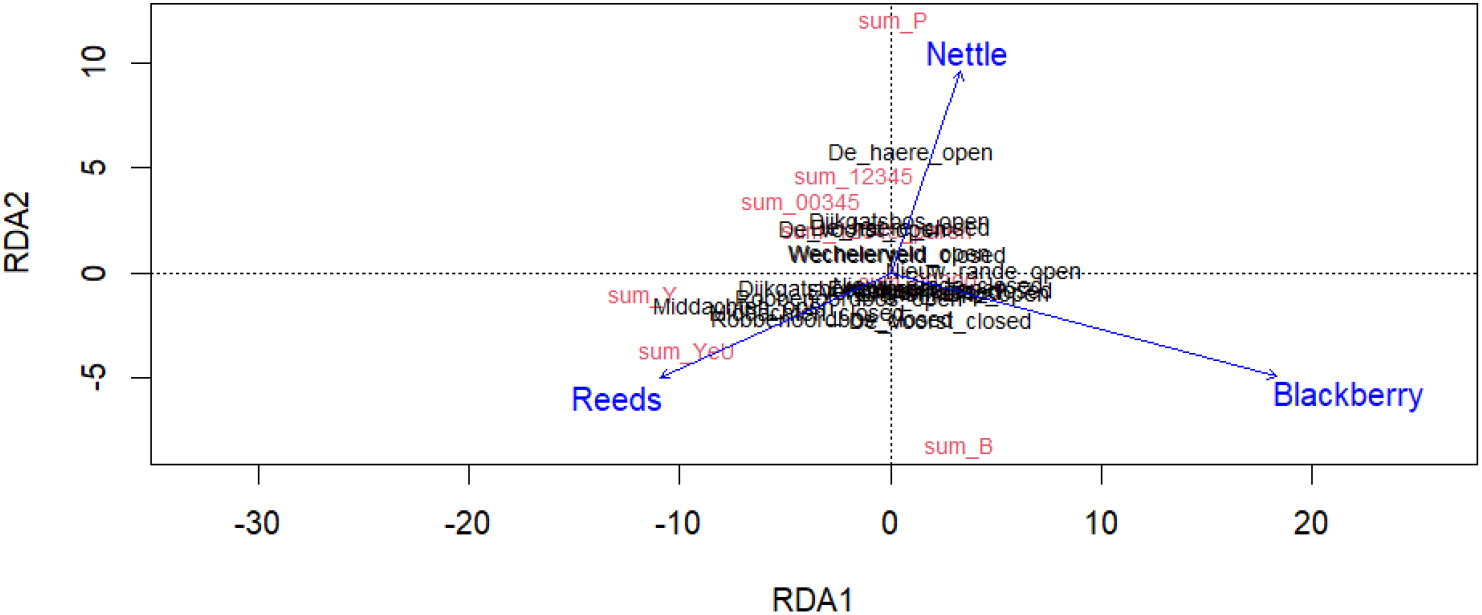
A multivariate analysis using three factors: nettle, reeds and blackberry dominances.

## Discussion

The aim of this research was to investigate the rate of evolutionary divergence of colour polymorphism of *Cepaea nemoralis* across different habitats of different age categories, especially when it comes to the time it takes for a population to establish evolutionary equilibrium of colour polymorphism. The expectation was that divergence would reach this equilibrium after approximately 300 years.

Congruent with Schilthuizen (2013), we found that the proportions of pink morphs tended to be higher in open areas. However, for the other morphs, no clear differences between open and closed habitats nor across age categories were found.

Furthermore, we resampled two locations previously also sampled by Schilthuizen (2013) for the following reason. If we were to find similar results more than a decade later, it would allow for both datasets to be combined. Of course, an additional reason was to see whether the populations sampled in 2011 had changed over time. Out of the four sample locations, only Robbenoordbos (closed) gave different results. Dijkgatsbos (open) also showed some discrepancies in the morph frequencies but its darkness score remained unchanged. Overall, this shows replicability of results, even 14 years later.

Contrary to the actual morph frequencies, the darkness scores do seem to give a more clear picture. The expectation was that when the different morphs in a population approach equilibrium, the difference in darkness score until at some point in time equilibrium would be reached. Given the significant differences within most habitat pairs and the increase in this difference over time that we found, our results do suggest ongoing evolutionary divergence at the temporal scale of our study. However, differences were small, and variation high, preventing any clear indication as to where a point of equilibrium might lie. Overall, it can be said that there is a tendency towards evolutionary equilibrium, although other effects that begin to play a role over longer periods of time might disturb a clear relation between darkness and age.

One of these effects might be an overlooked aspect that is currently clouding the results. We found a surprisingly large number of dark-coloured individuals in the open parts of the habitat pairs. These dark-shelled snails were mainly found in blackberry bushes, where they appeared (to a human observer) to be better camouflaged on the dark bramble branches.

Apart from temperature being a very important form of selection in determining shell morphology, it is also known that predation has a significant effect. Normally, these two selection pressures are expected to drive a population in the same direction: lighter morphs in an open area, and darker individuals when there is more shade. However, blackberry bushes do fall into the category of open habitats, but might favour dark individuals due to better camouflage. The fact is that the darkness score only factors in temperature, leaving camouflage out of the picture. The multivariate analysis does support this possibility, as blackberry bushes cover are able to explain darker morphs, while reeds cover explain light coloured morphs. Therefore, it might be wise in future studies to also include a camouflage score and combine it with the darkness score to obtain a more complete model.

A possible aid in such a future line of inquiry might be colour theory. A good way to quantify colour is Munsell’s (1905) colour system. This system defines colour in three aspects: value, hue, and chroma. In short, value is the degree of darkness of a colour, hue is the type of colour (for example yellow, red or green), and chroma is a measurement for the colour’s purity (Cochrane, S., 2014). In other words, every colour is defined and can be described by these three colour characteristics. This colour system could prove to be useful in designing a potential camouflage score, and we aim to develop this procedure in future papers on this subject.

## Supporting information

R script 1

