## Supplementary material for "Snails show their true colours: rate of evolutionary divergence in the shell colour polymorphism of *Cepaea nemoralis*": R script 1

R\_Script\_Cepaea\_GK


### R\_Script\_Cepaea\_GK

#### 2026-09-04

#### R Markdown

R script containing all analysis.

```{rm(list = ls())}

library(readxl) library(tidyverse) library(cowplot) library(dplyr)
library(stringr) library(tidyr) library(purrr) library(broom)
library(ape) library(phytools)

d <- read\_xlsx(“Results\_Govert\_Kwarten.xlsx”,1) str(d) d\(Cepaea\_type <- factor(d\)Cepaea\_type)
d\(Locality <- factor(d\)Locality)
d\(Habitat <- factor(d\)Habitat)
view(d)

# Proportion table.

YeU\_list <- c(“Y00000”, “Y00300”, “Y00345”, “Y003(45)”,
“Y00(34)5”, “Y00(345)”)

d\_flags <- d %>% mutate( digits\_only = gsub(“[^0-9]”, ““,
Cepaea\_type), is\_Y = str\_detect(Cepaea\_type,”^Y”), is\_B =
str\_detect(Cepaea\_type, “^B”), is\_P = str\_detect(Cepaea\_type, “^P”),
is\_00300 = str\_detect(Cepaea\_type, “00300”), is\_00345 =
str\_detect(Cepaea\_type, “00345”), is\_12345 = digits\_only == “12345”,
is\_12345\_paren = (digits\_only == “12345”) & str\_detect(Cepaea\_type,
“\([0-9]{2,}\)”), is\_YeU = Cepaea\_type %in% YeU\_list, is\_00000 =
str\_detect(Cepaea\_type, “^1^00000$”)  
)

# — Total N per locality —

totalN\_all <- d\_flags %>% group\_by(Locality) %>%
summarise(totalN\_all = sum(N, na.rm = TRUE), .groups = “drop”)

# — Total N excluding unbanded morphs —

totalN\_non00000 <- d\_flags %>% filter(!is\_00000) %>%
group\_by(Locality) %>% summarise(totalN\_non00000 = sum(N, na.rm =
TRUE), .groups = “drop”)

# — Total N of fivebanded morphs (with potential fusions) —

totalN\_12345 <- d\_flags %>% filter(is\_12345) %>%
group\_by(Locality) %>% summarise(total\_12345 = sum(N, na.rm = TRUE),
.groups = “drop”)

# — Totals for each group —

group\_sums <- d\_flags %>% group\_by(Locality) %>% summarise(
sum\_Y = sum(N \* is\_Y, na.rm = TRUE), sum\_B = sum(N \* is\_B, na.rm =
TRUE), sum\_P = sum(N \* is\_P, na.rm = TRUE), sum\_00300 = sum(N \*
is\_00300, na.rm = TRUE), sum\_00345 = sum(N \* is\_00345, na.rm = TRUE),
sum\_12345 = sum(N \* is\_12345, na.rm = TRUE), sum\_12345\_paren = sum(N \*
is\_12345\_paren, na.rm = TRUE), sum\_YeU = sum(N \* is\_YeU, na.rm = TRUE),
.groups = “drop” )

# — Combine denominators and calculate proportions —

prop\_table <- group\_sums %>% left\_join(totalN\_all, by =
“Locality”) %>% left\_join(totalN\_non00000, by = “Locality”) %>%
left\_join(totalN\_12345, by = “Locality”) %>% mutate( prop\_Y = sum\_Y /
totalN\_all, prop\_B = sum\_B / totalN\_all, prop\_P = sum\_P / totalN\_all,
prop\_YeU = sum\_YeU / totalN\_all,

```
prop_00300 = ifelse(totalN_non00000 > 0, sum_00300 / totalN_non00000, NA),
prop_00345 = ifelse(totalN_non00000 > 0, sum_00345 / totalN_non00000, NA),
prop_12345 = ifelse(totalN_non00000 > 0, sum_12345 / totalN_non00000, NA),

prop_12345_paren = ifelse(!is.na(total_12345) & total_12345 > 0,
                          sum_12345_paren / total_12345,
                          NA_real_)
```

) %>% select(Locality, totalN\_all, totalN\_non00000,
starts\_with(“prop\_”))

prop\_table view(prop\_table)

#Morph Proportions and Chi-square tests (including contigency table).
#——————————————————————————-

YeU\_list <- c(“Y00000”, “Y00300”, “Y00345”, “Y003(45)”,
“Y00(34)5”, “Y00(345)”)

d\_flags <- d %>% mutate( digits\_only = gsub(“[^0-9]”, ““,
Cepaea\_type), is\_Y = str\_detect(Cepaea\_type,”^Y”), is\_B =
str\_detect(Cepaea\_type, “^B”), is\_P = str\_detect(Cepaea\_type, “^P”),
is\_00300 = str\_detect(Cepaea\_type, “00300”), is\_00345 =
str\_detect(Cepaea\_type, “00345”), is\_12345 = digits\_only == “12345”,
is\_12345\_paren = (digits\_only == “12345”) & str\_detect(Cepaea\_type,
“\([0-9]{2,}\)”), is\_YeU = Cepaea\_type %in% YeU\_list, is\_not\_00000 =
!str\_detect(Cepaea\_type, “^2^00000$”) # exclude Y00000, B00000, P00000
)

# Total N per locality

totalN <- d\_flags %>% group\_by(Locality) %>%
summarise(totalN = sum(N, na.rm = TRUE), .groups = “drop”)

# Summed counts per morph group

group\_sums <- d\_flags %>% left\_join(totalN, by = “Locality”)
%>% group\_by(Locality, totalN) %>% summarise( sum\_Y = sum(N \*
is\_Y, na.rm = TRUE), sum\_B = sum(N \* is\_B, na.rm = TRUE), sum\_P = sum(N
\* is\_P, na.rm = TRUE), sum\_00300 = sum(N \* is\_00300, na.rm = TRUE),
sum\_00345 = sum(N \* is\_00345, na.rm = TRUE), sum\_12345 = sum(N \*
is\_12345, na.rm = TRUE), sum\_12345\_paren = sum(N \* is\_12345\_paren, na.rm
= TRUE), sum\_YeU = sum(N \* is\_YeU, na.rm = TRUE), .groups = “drop” )
%>% mutate( base = str\_remove(Locality, “\_(open|closed)\("),
habitat = ifelse(str\_detect(Locality, "open\)“),”open”,
“closed”) )

#view(group\_sums)

# Morphs to test

morph\_cols <- c(“sum\_Y”, “sum\_B”, “sum\_P”, “sum\_00300”,
“sum\_00345”, “sum\_12345”, “sum\_12345\_paren”, “sum\_YeU”)

chi\_for\_morph <- function(base\_name, morph, data) { d\_sub <-
data %>% filter(base == base\_name) if (nrow(d\_sub) != 2) return(NULL)
# Skip incomplete pairs

open <- d\_sub %>% filter(habitat == “open”) closed <- d\_sub
%>% filter(habitat == “closed”)

a <- open[[morph]] c <- closed[[morph]] total\_open <-
open\(totalN
total\_closed <- closed\)totalN

b <- total\_open - a d <- total\_closed - c

if (any(is.na(c(a, b, c, d))) || any(c(a, b, c, d) < 0))
return(NULL)

contingency <- matrix(c(a, b, c, d), nrow = 2, byrow = TRUE,
dimnames = list(c(“open”, “closed”), c(morph, “other”)))

test <- if (any(contingency < 5)) fisher.test(contingency) else
chisq.test(contingency) test\_type <- if (any(contingency < 5))
“Fisher” else “Chi-square”

contingency\_str <- paste( paste(rownames(contingency)[1], “:”,
paste(contingency[1, ], collapse = “,”)),
paste(rownames(contingency)[2], “:”, paste(contingency[2, ], collapse =
“,”)), sep = ” | ” )

tibble( habitat\_pair = base\_name, morph = morph, p\_value = test\(p.value,
statistic = unname(test\)statistic), df = if (“parameter”
%in% names(test)) unname(test$parameter) else NA, test\_type = test\_type,
contingency\_table = contingency\_str ) }

# Run all tests.

chi\_results\_all\_con <- map\_dfr(unique(group\_sums$base),
function(b) { map\_dfr(morph\_cols, function(m) chi\_for\_morph(b, m,
group\_sums)) })

chi\_results\_all\_con view(chi\_results\_all\_con)

#######################################CHI-SQUARE
TESTS########################## #Now for banded within habitats

prop\_table <- prop\_table %>% mutate( totalN\_all =
as.numeric(as.character(totalN\_all)), totalN\_non00000 =
as.numeric(as.character(totalN\_non00000)) )

site\_bases <- unique(sub(“\_(open|closed)(\_old|\_new)?\(", "",
prop\_table\)Locality))

chi\_banded\_test <- function(base, data) { open\_loc <-
paste0(base, “\_open”) closed\_loc <- paste0(base, “\_closed”)

d\_sub <- data %>% filter(Locality %in% c(open\_loc, closed\_loc))
%>% select(Locality, totalN\_all, totalN\_non00000) %>%
arrange(match(Locality, c(open\_loc, closed\_loc))) # ensure order: open,
closed if both present

if (nrow(d\_sub) != 2) {

```
return(tibble(
  base = base,
  open_loc = open_loc,
  closed_loc = closed_loc,
  note = "missing one or both localities",
  test_type = NA_character_,
  p_value = NA_real_,
  statistic = NA_real_,
  df = NA_real_,
  open_banded = NA_integer_,
  open_total = NA_integer_,
  closed_banded = NA_integer_,
  closed_total = NA_integer_,
  prop_open = NA_real_,
  prop_closed = NA_real_,
  prop_diff = NA_real_,
  ci_low = NA_real_,
  ci_high = NA_real_
))
```

}

# counts open\_banded <- d\_sub\(totalN\_non00000[1]
open\_total <- d\_sub\)totalN\_all[1] closed\_banded <-
d\_sub\(totalN\_non00000[2]
closed\_total <- d\_sub\)totalN\_all[2]

if (any(is.na(c(open\_banded, open\_total, closed\_banded,
closed\_total))) || any(c(open\_banded, open\_total, closed\_banded,
closed\_total) < 0)) { return(tibble( base = base, open\_loc =
open\_loc, closed\_loc = closed\_loc, note = “invalid counts/NA”, test\_type
= NA\_character\_, p\_value = NA\_real\_, statistic = NA\_real\_, df =
NA\_real\_, open\_banded = open\_banded, open\_total = open\_total,
closed\_banded = closed\_banded, closed\_total = closed\_total, prop\_open =
NA\_real\_, prop\_closed = NA\_real\_, prop\_diff = NA\_real\_, ci\_low =
NA\_real\_, ci\_high = NA\_real\_ )) }

# build contingency matrix: rows = groups (open, closed), cols =
banded/unbanded contingency <- matrix( c(open\_banded, open\_total -
open\_banded, closed\_banded, closed\_total - closed\_banded), nrow = 2,
byrow = TRUE, dimnames = list(c(open\_loc, closed\_loc), c(“Banded”,
“Unbanded”)) )

# choose test: Fisher if any expected < 5, else chi-square
use\_fisher <- any(chisq.test(contingency, simulate.p.value =
FALSE)\(expected < 5)
if (use\_fisher) {
test <- fisher.test(contingency)
test\_type <- "Fisher"
statistic <- NA\_real\_
df <- NA\_real\_
} else {
test <- chisq.test(contingency)
test\_type <- "Chi-square"
statistic <- unname(test\)statistic) df <-
unname(test$parameter) }

prop\_res <- prop.test(x = c(open\_banded, closed\_banded), n =
c(open\_total, closed\_total), correct = FALSE) prop\_diff <-
prop\_res\(estimate[1] -
prop\_res\)estimate[2] ci\_low <- prop\_res\(conf.int[1]
ci\_high <- prop\_res\)conf.int[2]

tibble( base = base, open\_loc = open\_loc, closed\_loc = closed\_loc,
note = NA\_character\_, test\_type = test\_type, p\_value = test\(p.value,
statistic = statistic,
df = df,
open\_banded = open\_banded,
open\_total = open\_total,
closed\_banded = closed\_banded,
closed\_total = closed\_total,
prop\_open = prop\_res\)estimate[1], prop\_closed =
prop\_res$estimate[2], prop\_diff = prop\_diff, ci\_low = ci\_low, ci\_high =
ci\_high ) }

# run across bases

chi\_banded\_results <- map\_dfr(site\_bases, chi\_banded\_test, data =
prop\_table)

# show results

chi\_banded\_results view(chi\_banded\_results)

#———————– MULTIVARIATE ANALYSIS———————————-
################################################################################
rm(list = ls())

d <- read\_xlsx(“Results\_Govert\_Kwarten.xlsx”,1) str(d) d\(Cepaea\_type <- factor(d\)Cepaea\_type)
d\(Locality <- factor(d\)Locality)
d\(Habitat <- factor(d\)Habitat)

library(dplyr) library(vegan)

env\_data <- read\_xlsx(“Results\_Govert\_Kwarten.xlsx”,23)
#view(env\_data) env\_data\(Locality <-
factor(env\_data\)Locality)

YeU\_list <- c(“Y00000”, “Y00300”, “Y00345”, “Y003(45)”,
“Y00(34)5”, “Y00(345)”)

# Assume your dataset has: Locality, Type, N

d\_flags <- d %>% mutate( digits\_only = gsub(“[^0-9]”, ““,
Cepaea\_type), is\_Y = str\_detect(Cepaea\_type,”^Y”), is\_B =
str\_detect(Cepaea\_type, “^B”), is\_P = str\_detect(Cepaea\_type, “^P”),
is\_00300 = str\_detect(Cepaea\_type, “00300”), is\_00345 =
str\_detect(Cepaea\_type, “00345”), # 12345 (with or without parentheses)
is\_12345 = digits\_only == “12345”, # 12345 with at least one set of
parentheses (around ≥2 digits) is\_12345\_paren = (digits\_only == “12345”)
& str\_detect(Cepaea\_type, “\([0-9]{2,}\)”), is\_YeU = Cepaea\_type
%in% YeU\_list, is\_00000 = str\_detect(Cepaea\_type, “^3^00000$”) # exact
matches to exclude )

# — Total N per locality —

totalN\_all <- d\_flags %>% group\_by(Locality) %>%
summarise(totalN\_all = sum(N, na.rm = TRUE), .groups = “drop”)

# — Total N excluding 00000 morphs —

totalN\_non00000 <- d\_flags %>% filter(!is\_00000) %>%
group\_by(Locality) %>% summarise(totalN\_non00000 = sum(N, na.rm =
TRUE), .groups = “drop”)

# — Total N of 12345 (with or without fusions) —

totalN\_12345 <- d\_flags %>% filter(is\_12345) %>%
group\_by(Locality) %>% summarise(total\_12345 = sum(N, na.rm = TRUE),
.groups = “drop”)

# — Compute sums for each group —

group\_sums <- d\_flags %>% group\_by(Locality) %>% summarise(
sum\_Y = sum(N \* is\_Y, na.rm = TRUE), sum\_B = sum(N \* is\_B, na.rm =
TRUE), sum\_P = sum(N \* is\_P, na.rm = TRUE), sum\_00300 = sum(N \*
is\_00300, na.rm = TRUE), sum\_00345 = sum(N \* is\_00345, na.rm = TRUE),
sum\_12345 = sum(N \* is\_12345, na.rm = TRUE), sum\_12345\_paren = sum(N \*
is\_12345\_paren, na.rm = TRUE), sum\_YeU = sum(N \* is\_YeU, na.rm = TRUE),
.groups = “drop” )

# — Combine denominators and compute proportions —

prop\_table <- group\_sums %>% left\_join(totalN\_all, by =
“Locality”) %>% left\_join(totalN\_non00000, by = “Locality”) %>%
left\_join(totalN\_12345, by = “Locality”) %>% mutate( # proportions
based on total N (all types) prop\_Y = sum\_Y / totalN\_all, prop\_B = sum\_B
/ totalN\_all, prop\_P = sum\_P / totalN\_all, prop\_YeU = sum\_YeU /
totalN\_all,

```
### proportions based on total non-00000 types
prop_00300 = ifelse(totalN_non00000 > 0, sum_00300 / totalN_non00000, NA),
prop_00345 = ifelse(totalN_non00000 > 0, sum_00345 / totalN_non00000, NA),
prop_12345 = ifelse(totalN_non00000 > 0, sum_12345 / totalN_non00000, NA),

### proportion of 12345 with fusions
prop_12345_paren = ifelse(!is.na(total_12345) & total_12345 > 0,
                          sum_12345_paren / total_12345,
                          NA_real_)
```

) %>% select(Locality, totalN\_all, totalN\_non00000,
starts\_with(“prop\_”))

prop\_table #view(prop\_table)

#Now go for the multivariate analysis

analysis\_data <- prop\_table %>% inner\_join(env\_data, by =
“Locality”)

morph\_cols <- c(“prop\_Y”, “prop\_B”, “prop\_P”, “prop\_00300”,
“prop\_00345”, “prop\_12345”, “prop\_12345\_paren”, “prop\_YeU”)

morph\_mat <- analysis\_data %>% select(all\_of(morph\_cols))
%>% as.data.frame()

# explanatory variables (plants)

plant\_cols <- setdiff(names(env\_data), “Locality”) plant\_mat <-
analysis\_data %>% select(all\_of(plant\_cols)) %>%
as.data.frame()

#view(plant\_mat)

sum\_cols <- c(“sum\_Y”, “sum\_B”, “sum\_P”, “sum\_00300”, “sum\_00345”,
“sum\_12345”, “sum\_12345\_paren”, “sum\_YeU”)

# Clean morph count matrix

snail\_data <- group\_sums %>% select(Locality, all\_of(sum\_cols))
%>% as.data.frame()

# Set rownames to Locality

rownames(snail\_data) <- snail\_data$Locality

# remove the Locality column to help Vegan a little bit

snail\_data <- snail\_data %>% select(-Locality)

head(snail\_data)

library(vegan) rda\_result <- rda(snail\_data ~ Blackberry + Reeds +
Nettle, data = plant\_mat) summary(rda\_result) vif.cca(rda\_result)
plot(rda\_result, scaling = 1.5)

#——— Relative frequency differences plotted against habitat age ———-
################################################################################

library(tidyr) library(dplyr) library(ggplot2)

prop\_table <- prop\_table %>% mutate( habitat =
ifelse(grepl(“open”, Locality), “Open”, “Closed”), pair =
sub(“\_(open|closed)$“,”“, Locality) ) %>% left\_join(d %>%
select(Locality, Age), by =”Locality”)

#view(prop\_table)

morph\_cols <- c(“prop\_Y”, “prop\_P”, “prop\_B”, “prop\_00300”,
“prop\_00345”, “prop\_12345”, “prop\_12345\_paren”, “prop\_YeU”)

prop\_long <- prop\_table %>% pivot\_longer(cols =
all\_of(morph\_cols), names\_to = “Morph”, values\_to = “Proportion”)

morph\_diff <- prop\_long %>% select(pair, habitat, Morph,
Proportion, Age) %>% pivot\_wider( names\_from = habitat, values\_from =
Proportion, values\_fn = mean,  
values\_fill = 0 ) %>% mutate(diff = abs(Open - Closed))

ggplot(morph\_diff, aes(x = Age, y = diff, color = Morph)) +
geom\_point(size = 2) + geom\_hline(yintercept = 0, linetype = “dashed”,
color = “grey40”) + theme\_minimal(base\_size = 13) +
scale\_color\_brewer(palette = “Dark2”) + labs( title = “Difference in
morph proportions vs habitat age all data”, x = “Habitat age (yrs)”, y =
“Absolute difference in proportion”, )

ggplot(morph\_diff, aes(x = Age, y = diff, color = Morph)) +
geom\_point(size = 2) + geom\_hline(yintercept = 0, linetype = “dashed”,
color = “grey40”) + theme\_minimal(base\_size = 13) +
scale\_color\_brewer(palette = “Dark2”, labels = c(“Y”, “P”, “B”, “00300”,
“00345”, “12345”, “Fusion”, “YeU”) ) + labs( title = “Difference in
morph proportions vs habitat age all data”, x = “Habitat age (yrs)”, y =
“Absolute difference in proportion”, color = “Morph category” )

#Correlation coefficient Pearson, returns nothing of value.

#view(morph\_diff)

library(dplyr) library(lme4) library(lmerTest)

cor\_results <- morph\_diff %>% group\_by(Morph) %>% summarise(
cor\_test = list(cor.test(Age, diff, method = “pearson”)), .groups =
“drop” ) %>% mutate( r = map\_dbl(cor\_test, ~ .x\(estimate),
p\_value = map\_dbl(cor\_test, ~ .x\)p.value), conf\_low =
map\_dbl(cor\_test, ~ .x\(conf.int[1]),
conf\_high = map\_dbl(cor\_test, ~ .x\)conf.int[2]) ) %>%
select(Morph, r, conf\_low, conf\_high, p\_value)

cor\_results

overall\_cor <- cor.test(morph\_diff\(Age,
morph\_diff\)diff, method = “pearson”) overall\_cor

#No correlation.

#———————————-DARKNESS SCORES——————————
################################################################################
rm(list = ls())

library(readxl) library(tidyverse) library(cowplot) library(purrr)
library(broom) library(ggplot2) library(dplyr)

d <- read\_xlsx(“Results\_Govert\_Kwarten.xlsx”,1) str(d)

d\(Locality <-
factor(d\)Locality) d\(Cepaea\_type
<- factor(d\)Cepaea\_type) d\(Habitat
<- factor(d\)Habitat)

#Now I have Mean\_DS and N and Locality.

darkness\_loc <- d %>% group\_by(Locality) %>% summarise(
total\_DS = sum(Combined\_DS, na.rm = TRUE), total\_N = sum(N, na.rm =
TRUE), mean\_darkness = total\_DS / total\_N ) %>% ungroup() %>%
mutate( pair = sub(“\_(open|closed)$“,”“, Locality), habitat =
ifelse(grepl(”open”, Locality), “Open”, “Closed”) )

expanded\_data <- d %>% mutate( pair = sub(“\_(open|closed)$“,”“,
Locality), habitat = ifelse(grepl(”open”, Locality), “Open”, “Closed”) )
%>% uncount(weights = N) %>% # expands each morph by its count N
select(Locality, pair, habitat, Individual\_DS)

kruskal\_results <- expanded\_data %>% group\_by(pair) %>%
group\_modify(~ broom::tidy( kruskal.test(Individual\_DS ~ habitat, data =
.x) )) %>% ungroup()

kruskal\_results

ggplot(expanded\_data, aes(x = habitat, y = Individual\_DS, fill =
habitat)) + geom\_boxplot() + facet\_wrap(~ pair, scales = “free\_y”) +
theme\_minimal() + labs( title = “Darkness score comparison per habitat
pair”, y = “Individual darkness score”, x = “Habitat type” ) +
scale\_fill\_manual(values = c(“Open” = “goldenrod”, “Closed” =
“darkgreen”))

kruskal.test(Individual\_DS ~ habitat, data = expanded\_data)

###############################PLOT THE
DIFFERENCES############################

darkness\_diff <- darkness\_loc %>% select(pair, habitat,
mean\_darkness) %>% tidyr::pivot\_wider(names\_from = habitat,
values\_from = mean\_darkness) %>% mutate( diff = Closed - Open )

darkness\_diff\(pair <-
factor(darkness\_diff\)pair, levels = c(“Dijkgatsbos”,
“Wechelerveld”, “Robbenoordbos”, “De\_haere”, “Nieuw\_rande”,
“Everdingen”, “Middachten”, “Kranenkamp”, “De\_voorst”))

ggplot(darkness\_diff, aes(x = pair)) + geom\_hline(yintercept = 0,
linetype = “dashed”, color = “gray60”) + geom\_point(aes(y = 0, color =
“Open”), size = 5) + geom\_point(aes(y = diff, color = “Closed”), size =
5) + geom\_segment(aes(x = pair, xend = pair, y = 0, yend = diff), color
= “black”) + scale\_color\_manual(values = c(“Open” = “yellow”, “Closed” =
“darkgreen”)) + labs( title = “Relative Darkness Difference”, x =
“Locality”, y = “Relative Darkness Score”, color = “Habitat” ) +
theme\_minimal(base\_size = 14) + theme( axis.text.x = element\_text(angle
= 45, hjust = 1), panel.grid.minor = element\_blank() )

#For the timeline plot

Age\_data <- read\_xlsx(“Results\_Govert\_Kwarten.xlsx”,1)
str(Age\_data)

Age\_data\(Locality <-
factor(Age\_data\)Locality)

darkness\_diff <- darkness\_loc %>% select(pair, habitat,
mean\_darkness) %>% tidyr::pivot\_wider(names\_from = habitat,
values\_from = mean\_darkness) %>% mutate( diff = Closed - Open )

darkness\_diff\(pair <-
factor(darkness\_diff\)pair, levels = c(“Dijkgatsbos”,
“Wechelerveld”, “Robbenoordbos”, “De\_haere”, “Nieuw\_rande”,
“Everdingen”, “Middachten”, “Kranenkamp”, “De\_voorst”))

darkness\_time <- darkness\_loc %>% left\_join(d %>%
select(Locality, Age), by = “Locality”)

darkness\_diff <- darkness\_time %>% group\_by(pair, habitat, Age)
%>% # group duplicates summarise(mean\_darkness = mean(mean\_darkness,
na.rm = TRUE), .groups = “drop”) %>% pivot\_wider(names\_from =
habitat, values\_from = mean\_darkness) %>% mutate(diff = Closed -
Open) # now Open and Closed are numeric

ggplot(darkness\_diff, aes(x = Age, y = diff)) + geom\_point(size = 3,
color = “darkred”) + geom\_smooth(method = “lm”, se = TRUE, color =
“black”) + theme\_minimal(base\_size = 14) + labs( title = “Darkness
Scores vs. Age”, x = “Habitat Age (yrs)”, y = “Difference in Darkness”
)

lm\_diff <- lm(diff ~ Age, data = darkness\_diff)
summary(lm\_diff)

#—————————OLD-NEW COMPARISONS———————————
################################################################################

rm(list=ls()) ###Time to look if Wieringemeer data are similar to
each other. #Plan, first proportions: #Check darkness scores

#Proportions: library(dplyr) library(stringr) library(tidyr)

d <- read\_xlsx(“Results\_Govert\_Kwarten.xlsx”,2) str(d) d\(Cepaea\_type <- factor(d\)Cepaea\_type)
d\(Locality <- factor(d\)Locality)
d\(Habitat <- factor(d\)Habitat)

YeU\_list <- c(“Y00000”, “Y00300”, “Y00345”, “Y003(45)”,
“Y00(34)5”, “Y00(345)”)

# Locality, Type, N

d\_flags <- d %>% mutate( digits\_only = gsub(“[^0-9]”, ““,
Cepaea\_type), is\_Y = str\_detect(Cepaea\_type,”^Y”), is\_B =
str\_detect(Cepaea\_type, “^B”), is\_P = str\_detect(Cepaea\_type, “^P”),
is\_00300 = str\_detect(Cepaea\_type, “00300”), is\_00345 =
str\_detect(Cepaea\_type, “00345”), is\_12345 = digits\_only == “12345”,
is\_12345\_paren = (digits\_only == “12345”) & str\_detect(Cepaea\_type,
“\([0-9]{2,}\)”), is\_YeU = Cepaea\_type %in% YeU\_list, is\_00000 =
str\_detect(Cepaea\_type, “^4^00000$”)  
)

# — Total N per locality —

totalN\_all <- d\_flags %>% group\_by(Locality) %>%
summarise(totalN\_all = sum(N, na.rm = TRUE), .groups = “drop”)

# — Total N excluding 00000 morphs —

totalN\_non00000 <- d\_flags %>% filter(!is\_00000) %>%
group\_by(Locality) %>% summarise(totalN\_non00000 = sum(N, na.rm =
TRUE), .groups = “drop”)

# — Total N of 12345 (with or without fusion) —

totalN\_12345 <- d\_flags %>% filter(is\_12345) %>%
group\_by(Locality) %>% summarise(total\_12345 = sum(N, na.rm = TRUE),
.groups = “drop”)

# — Compute sums for each group —

group\_sums <- d\_flags %>% group\_by(Locality) %>% summarise(
sum\_Y = sum(N \* is\_Y, na.rm = TRUE), sum\_B = sum(N \* is\_B, na.rm =
TRUE), sum\_P = sum(N \* is\_P, na.rm = TRUE), sum\_00300 = sum(N \*
is\_00300, na.rm = TRUE), sum\_00345 = sum(N \* is\_00345, na.rm = TRUE),
sum\_12345 = sum(N \* is\_12345, na.rm = TRUE), sum\_12345\_paren = sum(N \*
is\_12345\_paren, na.rm = TRUE), sum\_YeU = sum(N \* is\_YeU, na.rm = TRUE),
.groups = “drop” )

# — Combine denominators and compute proportions —

prop\_table <- group\_sums %>% left\_join(totalN\_all, by =
“Locality”) %>% left\_join(totalN\_non00000, by = “Locality”) %>%
left\_join(totalN\_12345, by = “Locality”) %>% mutate(

```
prop_Y = sum_Y / totalN_all,
prop_B = sum_B / totalN_all,
prop_P = sum_P / totalN_all,
prop_YeU = sum_YeU / totalN_all,


prop_00300 = ifelse(totalN_non00000 > 0, sum_00300 / totalN_non00000, NA),
prop_00345 = ifelse(totalN_non00000 > 0, sum_00345 / totalN_non00000, NA),
prop_12345 = ifelse(totalN_non00000 > 0, sum_12345 / totalN_non00000, NA),


prop_12345_paren = ifelse(!is.na(total_12345) & total_12345 > 0,
                          sum_12345_paren / total_12345,
                          NA_real_)
```

) %>% select(Locality, totalN\_all, totalN\_non00000,
starts\_with(“prop\_”))

#view(prop\_table)

#————Chi-square tests between open/old of the same habitat————–
################################################################################
#same as before library(dplyr) library(stringr) library(purrr)
library(broom)

YeU\_list <- c(“Y00000”, “Y00300”, “Y00345”, “Y003(45)”,
“Y00(34)5”, “Y00(345)”)

d\_flags <- d %>% mutate( digits\_only = gsub(“[^0-9]”, ““,
Cepaea\_type), is\_Y = str\_detect(Cepaea\_type,”^Y”), is\_B =
str\_detect(Cepaea\_type, “^B”), is\_P = str\_detect(Cepaea\_type, “^P”),
is\_00300 = str\_detect(Cepaea\_type, “00300”), is\_00345 =
str\_detect(Cepaea\_type, “00345”), is\_12345 = digits\_only == “12345”,
is\_12345\_paren = (digits\_only == “12345”) & str\_detect(Cepaea\_type,
“\([0-9]{2,}\)”), is\_YeU = Cepaea\_type %in% YeU\_list, is\_not\_00000 =
!str\_detect(Cepaea\_type, “^5^00000$”) # exclude Y00000, B00000, P00000
)

totalN <- d\_flags %>% group\_by(Locality) %>%
summarise(totalN = sum(N, na.rm = TRUE), .groups = “drop”)

group\_sums <- d\_flags %>% left\_join(totalN, by = “Locality”)
%>% group\_by(Locality, totalN) %>% summarise( sum\_Y = sum(N \*
is\_Y), sum\_B = sum(N \* is\_B), sum\_P = sum(N \* is\_P), sum\_00300 = sum(N \*
is\_00300), sum\_00345 = sum(N \* is\_00345), sum\_12345 = sum(N \* is\_12345),
sum\_12345\_paren = sum(N \* is\_12345\_paren), sum\_YeU = sum(N \* is\_YeU),
.groups = “drop” ) %>% mutate( site\_habitat = str\_remove(Locality,
“\_(old|new)\("),
period = ifelse(str\_detect(Locality, "old\)“),”old”,
“new”) )

morph\_cols <- c(“sum\_Y”, “sum\_B”, “sum\_P”, “sum\_00300”,
“sum\_00345”, “sum\_12345”, “sum\_12345\_paren”, “sum\_YeU”)

chi\_for\_morph\_time <- function(site\_name, morph, data) { d\_sub
<- data %>% filter(site\_habitat == site\_name) if (nrow(d\_sub) !=
2) return(NULL) # skip incomplete sites

old <- d\_sub %>% filter(period == “old”) new <- d\_sub %>%
filter(period == “new”)

a <- old[[morph]] c <- new[[morph]] total\_old <- old\(totalN
total\_new <- new\)totalN

b <- total\_old - a d <- total\_new - c

if (any(is.na(c(a,b,c,d))) || any(c(a,b,c,d) < 0))
return(NULL)

contingency <- matrix(c(a,b,c,d), nrow = 2, byrow = TRUE, dimnames
= list(c(“old”, “new”), c(morph, “other”)))

test <- if (any(contingency < 5)) fisher.test(contingency) else
chisq.test(contingency)

tibble( site\_habitat = site\_name, morph = morph, p\_value = test\(p.value,
statistic = test\)statistic, df = if (“parameter” %in%
names(test)) test$parameter else NA, test\_type = if (any(contingency
< 5)) “Fisher” else “Chi-square” ) }

# Run for all sites and all morphs

chi\_results\_time <- map\_dfr(unique(group\_sums$site\_habitat),
function(b) { map\_dfr(morph\_cols, function(m) chi\_for\_morph\_time(b, m,
group\_sums)) })

view(chi\_results\_time) #Ok, most are not significant, however still
some are… #The significant ones:

chi\_results\_time %>% mutate(significant = p\_value < 0.05)
%>% count(site\_habitat, significant)

#let’s make the diagram as well:

morph\_cols <- c(“prop\_Y”, “prop\_B”, “prop\_P”, “prop\_00300”,
“prop\_00345”, “prop\_12345”, “prop\_12345\_paren”, “prop\_YeU”)

# Prepare the numeric matrix

prop\_numeric <- prop\_table %>% select(Locality,
all\_of(morph\_cols)) %>% mutate(across(all\_of(morph\_cols),
as.numeric)) %>% tibble::column\_to\_rownames(“Locality”)

# Compute Euclidean distances

dist\_matrix <- dist(prop\_numeric, method = “euclidean”)

#Ward method hc\_ward <- hclust(dist\_matrix, method =
“ward.D2”)

plot(hc\_ward, main = “Phenogram Old-New Comparison”, xlab =
“Localities”, ylab = “Height (Dissimilarity)”, sub = ““, cex = 0.8)

#Also here, Robbenoordbos closed results are not as expected.

#Let’s check the darkness scores:

d\_dark <- d %>% mutate( base =
sub(“*(open|closed)*(old|new)\(",
"", Locality),
habitat = ifelse(grepl("open", Locality),
"open", "closed"),
period = ifelse(grepl("old\)”, Locality), “old”, “new”)
)

expanded\_dark <- d\_dark %>% uncount(weights = N) %>%
select(base, habitat, period, Individual\_DS)

kruskal\_oldnew <- expanded\_dark %>% group\_by(base, habitat)
%>% group\_modify(~ { if (length(unique(.x$period)) > 1) {
broom::tidy(kruskal.test(Individual\_DS ~ period, data = .x)) } else {
tibble(statistic = NA, p.value = NA) } }) %>% ungroup()

print(kruskal\_oldnew)

```

---

1. YBP↩︎
2. YBP↩︎
3. YBP↩︎
4. YBP↩︎
5. YBP↩︎
